# Inflammation induces pro-NETotic neutrophils via TNFR2 signaling

**DOI:** 10.1101/2021.06.21.448937

**Authors:** Friederike Neuenfeldt, Jan Christoph Schumacher, Ricardo Grieshaber-Bouyer, Jüri Habicht, Jutta Schröder-Braunstein, Annika Gauss, Beate Niesler, Niko Heineken, Alexander Dalpke, Matthias M. Gaida, Thomas Giese, Stefan Meuer, Yvonne Samstag, Guido Wabnitz

**Author notes:** FN and JCS contributed equally to this manuscript.

## Abstract

Cytokines released during chronic inflammatory diseases induce pro-inflammatory properties in polymorphonuclear neutrophils (PMN). Here we show that *in vitro* cytokine treatment leads to the development of a subgroup of human PMN expressing CCR5, termed CCR5^+^ cytokine-induced PMN (CCR5^+^ cPMN). Auto/paracrine TNF signaling increases intracellular neutrophil elastase (ELANE) abundance and induces NETosis in CCR5^+^ cPMN. Triggering of CCR5 amplifies NETosis. Membranous TNF (mTNF) outside-in signaling induces the formation of reactive oxygen species, a known activator of NETosis. *In vivo*, we find an increased number of CCR5^+^ cPMN in the peripheral blood and inflamed lamina propria of patients with ulcerative colitis (UC) but not Crohn’s disease (CD). Notably, failure of anti-TNF therapy is associated with higher frequencies of CCR5^+^ cPMN. In conclusion, we identify a phenotype of pro-NETotic, CCR5 positive PMN present in inflamed tissue *in vivo* and inducible *in vitro*. These cells may reflect an important component of tissue damage during chronic inflammation and could be of diagnostic value.

## Introduction

Polymorphonuclear neutrophils (PMN) represent the first line of defense against invading pathogens and are involved in inflammatory processes. PMN kill bacteria by releasing anti-bacterial proteins (degranulation), production of reactive oxygen species (ROS; oxidative burst) and by phagocytosis. Another important antibacterial mechanism is the formation of neutrophil extracellular traps (NETs). NET formation – also termed NETosis – is a process in which neutrophils discharge chromatin, forming an extracellular mesh of DNA, oxidases, and proteases, e.g. neutrophil elastase (ELANE). NETs are believed to capture and kill pathogens as well as to present them to other phagocytes on a large scale (*1*). However, NETs can also initiate destructive processes through e.g. interaction with other leukocytes. Consequently, exaggerated NETosis or a failure in the removal of NETs by macrophages was shown to play an important role in sustaining or exacerbating immune-mediated inflammatory diseases as well as cancer (*2–4*).

Pro-inflammatory cytokines such as TNF occurring in inflamed tissues can induce NETosis (*5*). On the other hand, the lifespan of PMN is strongly increased by pro-inflammatory cytokines like IFNγ or GM-CSF (*6–9*). Whether PMN die by NETosis or survive by delaying apoptosis is an important decision point for progression of inflammatory diseases. NETosis regulation has therefore moved into the focus of clinical research. In that regard, one hypothesis claims that PMN subsets exist exhibiting variable sensitivities to NETosis (*10*). Today it is known that PMN heterogeneity exists due to circadian rhythm or homeostatic turnover (*11–13*) and functionally or phenotypically altered PMN have been described in tumors, infections and in inflamed tissues (*14–19*). It is currently unknown whether PMN diversification with an altered functional repertoire or susceptibility to NETosis could explain increased NETosis in inflammed tissues.

In this study, we analyzed functional and phenotypical alterations of human PMN under inflammatory conditions. GM-CSF and IFNγ were used to prime PMN, prolong their survival and allow (trans)differentiation as described before for *in vitro* and *in vivo* experiments (*20–23*). A gene expression analysis was performed with freshly isolated human PMN and cytokine-stimulated PMN (cPMN). We discovered a cytokine-induced diversification of human PMN and significant upregulation of CCR5 on one cytokine-depended subgroup, termed CCR5^+^ cytokine-depended (c)PMN.

## Results

### Cytokine stimulation induces a CCR5-positive PMN subset

We began by examining gene expression changes induced by extended cytokine stimulation of human neutrophils. We purified resting whole blood PMN (rPMN) from healthy volunteers by density centrifugation followed by magnetic bead isolation (Fig. 1A), yielding 99% pure CD66b^+^ rPMN without eosinophil contamination (Supplementary Figure 1A). rPMN were cultured with GM-CSF plus IFNγ for 48 hours to obtain cytokine-induced PMN (cPMN) as described previously (*20–23*). As expected, culture of PMN without cytokines led to a higher rate of apoptosis and therefore further experiments were performed with cytokine-induced PMN (Supplementary Figure 1B).

**Figure 1.**
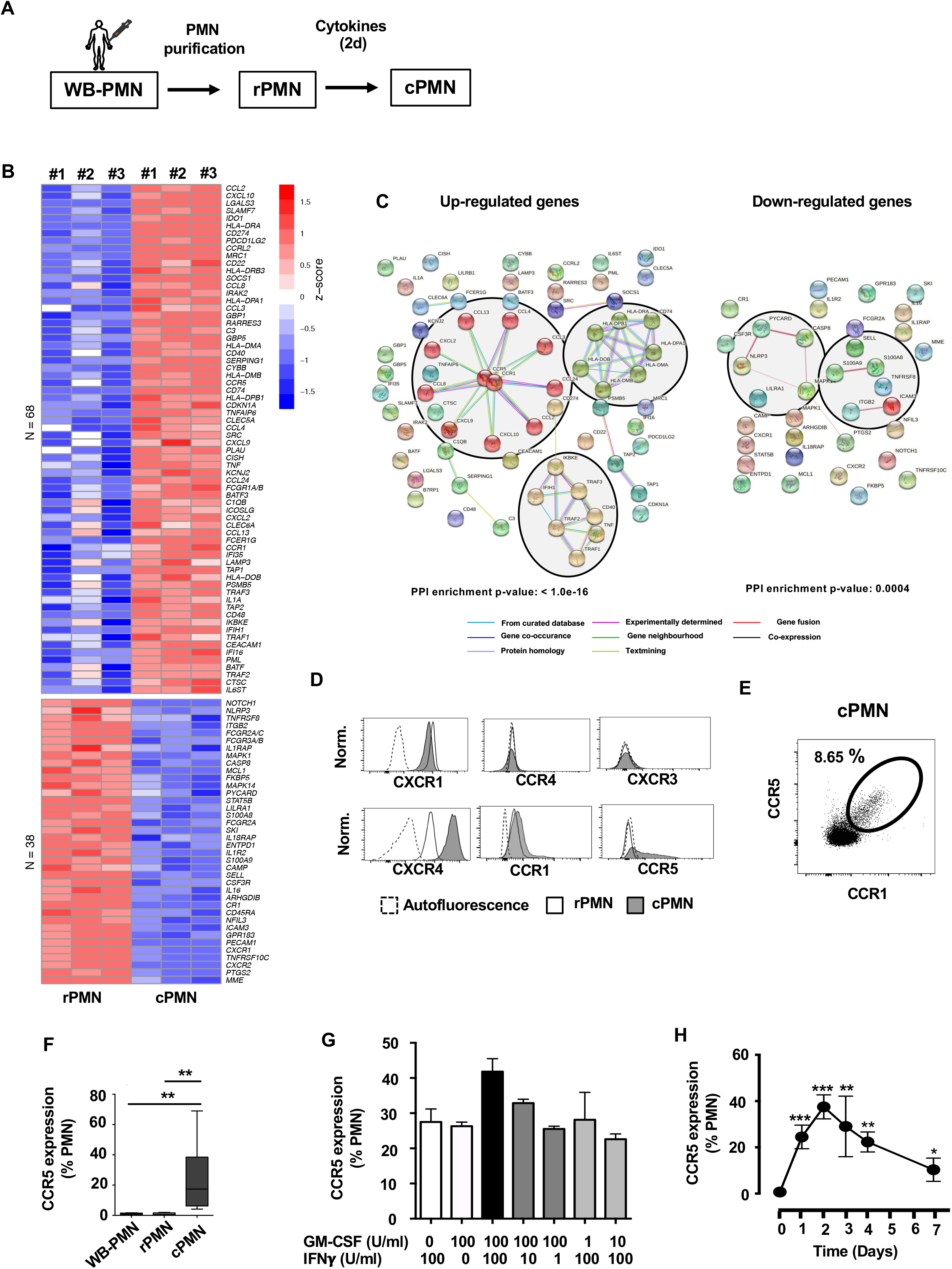
Cytokine stimulation of human PMN induces the development of a CCR5-expressing PMN subgroup (CCR5^+^ cPMN) **A)** Whole blood PMN (WB-PMN) from healthy individuals were purified by density gradient centrifugation followed by negative magnetic bead isolation obtaining resting PMN (rPMN). These cells were activated by the cytokines GM-CSF (100 U/ml) and IFNγ (10 ng/ml) for two days resulting in cytokine-incubated PMN (cPMN) that were used for RNA isolation. **B)** Heatmap of NanoString nCounter-based (NanoString Technologies, Inc.) comparing rPMN and cPMN of three different blood donors. Only genes with an expression value of ≥ 100 CodeSet counts in all three replicates in one group were considered. **C)** Interaction network among up (left) and down (right) regulated genes found in the nCounter analysis of rPMN and cPMN. The graph was generated from STRING database. Connections reflect physical protein interactions. The determination of the connection is shown below as color code. A small protein-protein interaction (PPI) enrichment p-value indicate that the nodes are not random and that the observed number of edges is significant. **D)** The surface expression of chemokine receptors in rPMN (open histograms) and cPMN (tinted histograms) was analyzed using flow cytometry. The autofluorescence of cPMN is shown as dotted histogram. The histograms are representative for three independent experiments. **E-F)** CCR1 and CCR5 co-expression was analyzed on cPMN by (double-staining) flow cytometry (**E**, dot plot). The amount of CCR5^+^ cPMN of total PMN was quantified and is depicted for 20 independent experiments in **F** (n=20; ANOVA; **p<0.01). **G)** The percentage of CCR5^+^ cPMN was assessed after two days of incubation of primary human PMN with the indicated concentrations of GM-CSF and IFNγ (n=4). **H)** Percentage of CCR5^+^ cPMN was quantified at day 0 (3h after adding cytokines) and at days 1-3 using flow cytometry. Statistics represent ANOVA test with d0 as control value (SEM; n=4; **p<0.05; ***p<0.01).

We identified 106 differentially expressed genes between rPMN and cPMN at log_2_ fold change ≥ 2 and a false discovery rate of 0.01. 38 genes were downregulated in cPMN and 68 genes were significantly up-regulated (Fig. 1B). *CXCR2* was downregulated in stimulated PMN. Moreover, *NLRP3, CASP8* and *PYCARD*, three transcripts related to the NLRP3 inflammasome-complex were downregulated. We also found downregulation of transcripts related to bacterial defense including *CAMP* (encoding cathelicidin) and CR1 (encoding complement receptor 1). Transcripts connected to apoptosis or PMN removal (*CASP8* and *PECAM1*), adhesion molecules (*SELL* and *ITGB2*) and phagocytosis (*ICAM3*, *PECAM1* and *ITGB2*) were also downregulated.

Among the upregulated transcripts were chemokines (*CCL2*, *CCL3*, *CCL4*, *CCL8*, *CCL13*, *CXCL2*, *CXCL9*, *CXCL10* and *CCL24*) and chemokine receptors (*CCR1*, *CCR5*), the proinflammatory cytokine *TNF*, complement factors and complement regulators (*C1QB*, *C3*, *SERPING1*), a transcript encoding a catalytic subunit of the NADPH-oxidase (*CYBB*) and transcripts encoding proteins involved in antigen-presentation and lymphocyte regulation (*HLA-DMA*, *HLA-DMB*, *HLA-DRA*, *HLA-DPA1*, *HLA-DRB3*, *CD74*, *PSMB5*, *TAP1*, *TAP2*, *CD274*, *ICOSLG*, *CD48*).

We then used the STRING database (Search Tool for the Retrieval of Interacting *Genes*/Proteins) to analyze networks of predicted protein-protein interactions and pathways among the differentially expressed genes (*24*). In this quality-controlled association database only direct (physical) interactions were included. Most up- and downregulated genes were not connected. However, three closely connected clusters comprising chemokine signaling via CCR1 and CCR5, antigen-processing, and NF-kB signaling (including TNF) were among in the upregulated transcripts (Fig. 1C). Down-regulated transcripts clustered in NLRP3 inflammasome-complex-related genes and genes involved in adhesion and exocytosis.

Chemokine receptor expression was analyzed on protein level using flow cytometry. The chemokine receptors CCR4 and CXCR3 were not expressed in rPMN and cPMN. CXCR4 was expressed in low amounts on rPMN (CXCR4^dim^) and was strongly upregulated in cPMN (CXCR4^bright^) (Fig. 1D). CCR1 was already expressed in rPMN, but cytokine stimulation induced a bimodal expression pattern in cPMN. While CCR5 was hardly detectable on the surface of rPMN, its expression was moderately upregulated on cPMN. Notably, CCR5 was not upregulated on all cPMN, but again restricted to only a subgroup of cPMN. CCR1^bright^ and CCR5-positive cPMN marked the same subgroup (Fig. 1E). Thus, cytokine treatment induced two subgroups of cPMN characterized by simultaneous expression or absence of CCR5, termed CCR5^+^ cPMN and CCR5^−^ cPMN. We next examined the frequency of PMN phenotypes across conditions. Less than 1% of whole blood (WB)-PMN or rPMN expressed CCR5. The amount of CCR5^+^ cells in cPMN increased to 4 – 69% with a median of 17.4% after cytokine incubation (Fig. 1F). Titration experiments covering a range of GM-CSF and IFNγ observed in colitis models *in vivo* showed that CCR5 expression was dose-dependent and increased in particular with higher concentrations of INFγ (Fig. 1G) (*25–28*). Interestingly, both INFγ and GM-CSF alone were sufficient to induce CCR5^+^ cPMN, suggesting that this diversification process is not unique to either cytokine. CCR5 was not detected 3h after adding of the cytokines (day 0) and peaked after two days (Fig. 1H). Based on these experiments, 100 U/ml GM-CSF + 10 ng/ml IFNγ and an incubation time of 2 days were chosen as optimum parameters.

### *De novo* expression and surface transport of CCR5

Western blot analysis from whole cell lysates independently confirmed CCR5 expression (Fig. 2A). Unexpectedly, CCR5 was expressed in both rPMN and cPMN at comparable levels, suggesting intracellular storage of CCR5 and mobilization to the surface upon cytokine activation (Fig. 2A and B). In flow cytometry, freshly isolated PMN (rPMN) did not show surface expression of CCR5, however, strong intracellular expression was again recognizable in permeabilized cells (Fig. 2C). Similarly, cPMN contained high levels of intracellular CCR5 but only a subgroup of cPMN expressed CCR5 on the cell surface. Super-resolution microscopy using structured-illumination microscopy (3D-SIM) revealed that CCR5 was stored in vesicular-like structures in rPMN and cPMN, while surface localization was only detectable on a fraction of cPMN (Fig. 2D). A staining with fluorescently-labelled phalloidin and imaging flow cytometry showed that the surface expression of CCR5 was accompanied with lower F-actin content, while CRR5^−^ cPMN had an increased F-actin content (Fig. 2E). Accordingly, pertubation of actin dynamics with cytochalasin D (CytoD) led to higher surface expression of CCR5 and stabilization of F-actin with Jasplakinolide (Jas) diminished the amount of CCR5 on the cell surface (Fig. 2F).

**Figure 2.**
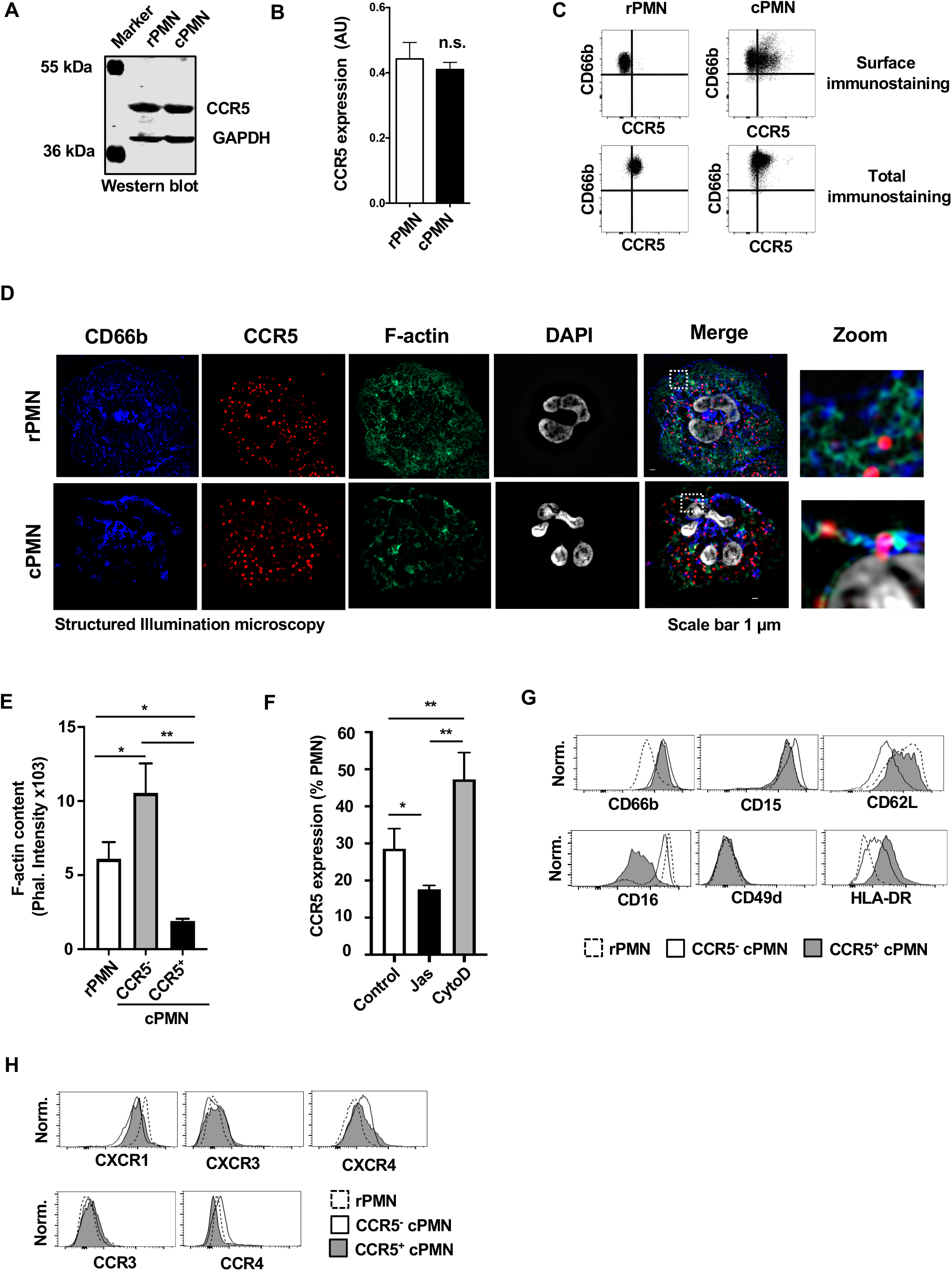
CCR5 is newly expressed and expelled from intracellular stores. **A-B)** Total CCR5 expression was detected in whole cell lysates from rPMN and cPMN by western blot analysis. GAPDH immunostaining served as loading control. The blot in A is representative for three independent experiments. A quantification of the CCR5 expression as ratio between CCR5- and GAPDH-derived signals is depicted in B (SEM; n=3; t-test; n.s.=not significant). **B)** CCR5 was detected on rPMN (left) and cPMN (right) by surface immunostaining (upper part) or by total immunostaining (lower part) using 0.1% saponin to permeabilize the cells. Shown are representative flow cytometry histograms taken from three independent experiments. **C)** rPMN and cPMN were adhered on coverslips, fixed, permeabilized and immunostained for CD66b (blue) and CCR5 (red). In addition F-actin (phalloidin-AF488) and nuclei (DAPI) were labelled. Images of 28 cells from three independent experiments were acquired using structured-illumination microscopy (3D-SIM). **D)** rPMN and cPMN were fixed and stained with phalloidin-AF488 to quantify F-actin by imaging flow cytometry. Shown is the total intensity from 6 independent experiments (n=6, SEM, ANOVA, *p<0.05, **p<0.01). **E)** PMN were stimulated with GM-CSF/ IFNγ in the presence or absence of either CytoD (250nM) or Jas (500nM) for one day. The expression of CCR5 was quantified using flow cytometry (n=3, SEM, ANOVA, *p<0.05, **p<0.01). **G-H)** The expression of PMN lineage-associated receptors (**G)**, CD66b, CD15, CD62L, CD16, HLA-DR and CD49d) as well as chemokine receptors (**H)**, CXCR1, CXCR3, CXCR4, CCR3, CCR4) were analyzed on rPMN, CCR5^−^ and CCR5^+^ cPMN by flow cytometry. The cells were gated for CD66b^+^ (rPMN) and their expression of CCR5 (CD66b^+^ cPMN). The histograms are representative for three independent experiments.

### Diversification-dependent regulation of surface receptors

We next used flow cytometry to phenotype rPMN, CCR5^−^ and CCR5^+^ cPMN using a selection of surface markers associated with neutrophil activation. Both CCR5^+^ and CCR5^−^ cPMN showed an activated phenotype with upregulation of CD66b (Fig. 2G), while CD15 moderately upregulated on CCR5^−^ but not CCR5^+^ cPMN. Further, CCR5^−^ cPMN were CD62L^dim^CD16^bright^ and CCR5^+^ cPMN were CD62L^bright^CD16^dim^. Notably, the gene encoding CD62L (SELL) was also downregulated on mRNA level in activated neutrophils (Fig. 1B).

The integrin CD49d (α-chain of the heterodimeric integrin VLA-4 (very-late-antigen-4) was not expressed in CCR5^−^ and CCR5^+^ cPMN (Fig. 2G). HLA-DR was not expressed on rPMN but was induced on both CCR5^+^ and CCR5^−^ cPMN, mirroring the upregulation of HLA-DRA on transcript level (Fig. 1C). The upregulation was stronger on CCR5^+^ cPMN (HLA-DR^bright^) compared to CCR5^−^ cPMN (HLA-DR^dim^). CXCR1, one of the major chemokine receptors expressed on PMN, was slightly downregulated on CCR5^−^ and CCR5^+^ cPMN compared to rPMN (Fig. 2H). An upregulation of CXCR4 could be observed on the surface of both CCR5^−^ and CCR5^+^ cPMN, with a higher expression on CCR5^−^ cPMN (CXCR4^bright^). No significant upregulation of CXCR3 or CCR3 was detectable on CCR5^−^ or CCR5^+^ cPMN. Taken together, the cytokines GM-CSF and IFNγinduce a diversification of PMN into CCR5^−^CXCR4^bright^HLA-DR^dim^CD62L^dim^CD16^brigh^ and CCR5^+^CXCR4^dim^HLA-DR^bright^CD62L^bright^CD16^dim^ cPMN.

### CCR5^+^ cPMN have a reduced migratory, exocytotic and phagocytic capacity

The expression of CCR5 and CXCR4 on cPMN suggests a homing potential of these cells towards inflamed sites. We used imaging flow cytometry to quantify the polarization of PMN, which can be associated with migratory behavior (*29*). Cells were termed polarized if they displayed formation of an F-actin rich protruding front (lamellipodium) and a retracting rear, named uropod. cPMN were incubated with the CXCR4-ligand SDF1α (100ng/ml) or the CCR5-ligand CCL5 (100ng/ml) for 5 min, fixed and stained for F-actin (phalloidin) and CCR5. While most cells had a round morphology in the absence of chemokines, a clear polarization of F-actin and concomitantly a lamellipodium formation could be observed for both CCR5^−^ and CCR5^+^ cPMN after SDF1α stimulation (Fig. 3A). In line with lower CXCR4 expression on CCR5^+^ cPMN, their polarization in response to SDF1 was weaker compared to CXCR4^bright^ CCR5^−^ cPMN (Fig. 3B). Conversely, CCL5 induced a polarization in CCR5^+^ but not CCR5^−^ cPMN (Fig. 3A and B). Thus, CCR5^−^ cPMN appear to be particularly responsive to CXCR4, while CCR5^+^ cPMN are more sensitive to CCL5.

**Figure 3.**
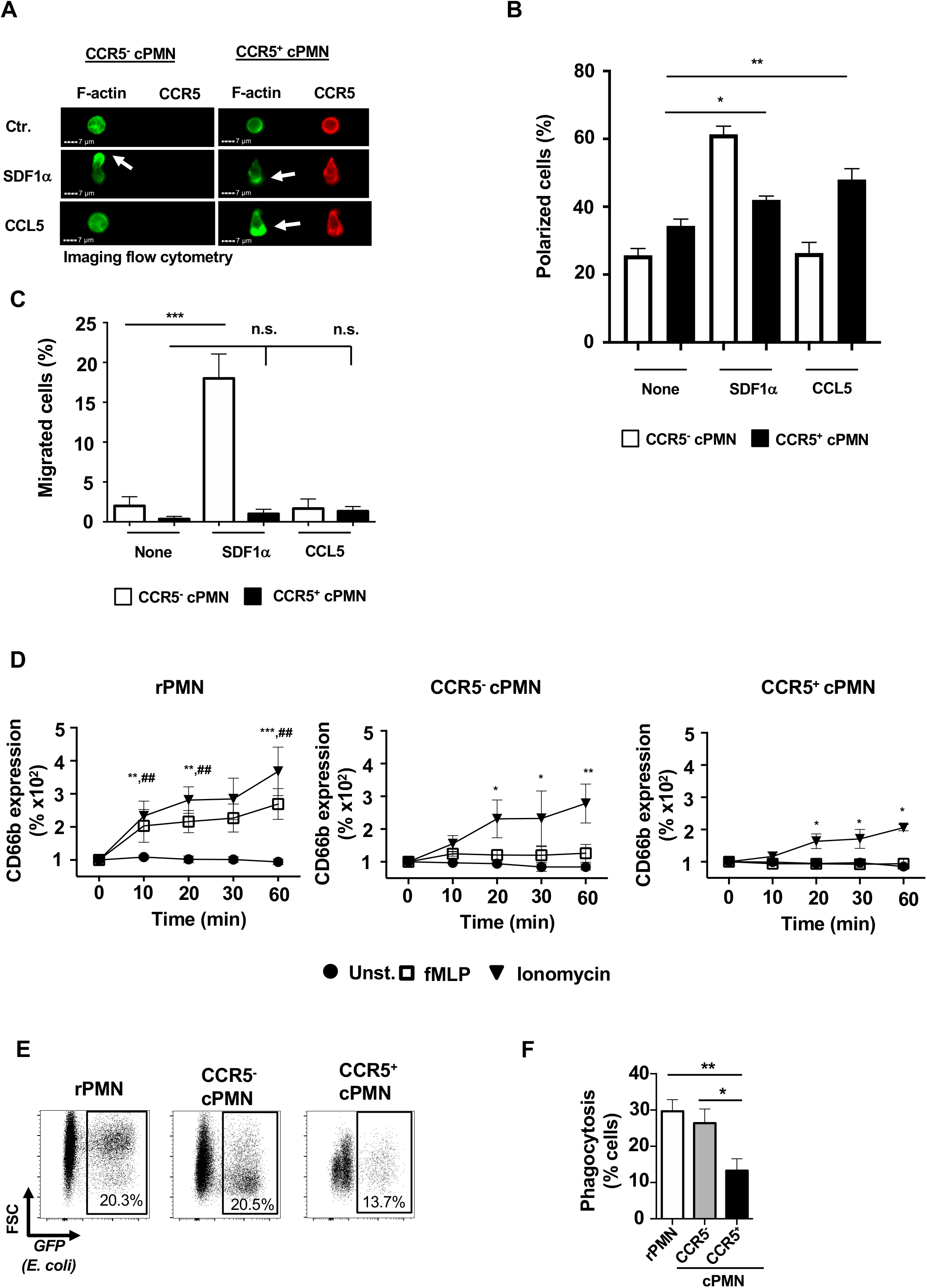
Diminished migration and bacterial defense mechanisms in CCR5^+^ cPMN. **A)** cPMN were stimulated with SDF1α (100ng/ml) or CCL5 (100ng/ml) for 5 minutes. Then, cells were fixed and stained for F-actin (green) and CCR5 (red). Images of 25,000 cells were acquired using imaging flow cytometry. Shown are representative images of CCR5^−^ cPMN (left) and CCR5^+^ cPMN (right) that were either left untreated (upper panel) or stimulated as indicated (lower panels). The arrows indicate polarized F-actin as it occurs after cell polarization. **B)** Quantification of the percentage of polarized CCR5^−^ (white bars) and CCR5^+^ cPMN (black bars) after treatment with solvent control (Ctr.), SDF1α or CCL5 is shown (n=3, SEM, ANOVA, *p<0.05, **p<0.01). **C)** Migration of PMN was performed in a transwell assay. Migrated cells were counted using flow cytometry and immunostaining for CCR5. Migrated cells are depicted as percent of the number of input cells of the respective subgroup (n=3, SEM, ANOVA, ***p<0.01, n.s.=not significant). **D)** rPMN or cPMN were stimulated for one hour with fMLP or ionomycin. Afterwards, the expression level of CD66b was measured for rPMN (left graph), CCR5^−^ cPMN (middle graph) and CCR5^+^ cPMN (right graph) by flow cytometry and normalized to time point zero of the respective kinetics. Shown are the percentage changes as mean value from 5 experiments (n=5; SEM; ANOVA with t=0 min as control; *p<0.05, **p<0.01 and ***p<0.001 of ionomycin; ## p<0.01 of fMLP). **E-F)** Phagocytosis of *E. coli* was analyzed for rPMN, CCR5^−^ cPMN and CCR5^+^ cPMN using flow cytometry. Sample dot plots are shown in **E** and a quantification of PMN that phagocytosed *E. coli* is shown in **F** (n=5; ANOVA, SEM, *p<0.05; **p<0.01).

To test if polarization by SDF1α or CCL5 also induces migration, we analyzed the migratory behavior of cPMN in a transwell assay over three hours (Fig. 3C). Concordant with polarization results from imaging flow cytometry, SDF1α, but not CCL5 induced migration of CXCR4^bright^ CCR5^−^ cPMN. Interestingly, we did not observe migration of CXCR4^dim^ CCR5^+^ cPMN in response to either CCL5 or SDF1α, suggesting that CCR5^+^ cPMN develop in inflammatory conditions, but do not migrate into inflamed sites on large scale.

Next, we examined exocytosis and phagocytosis in rPMN and cPMN. Exocytosis was assessed by flow cytometry analysis of surface expression of CD66b, which is stored in the membrane of specific and tertiary granules and is transported to the cell surface upon PMN activation. Both fMLP and ionomycin induced rapid exocytosis of CD66b in rPMN starting after 10 min and reaching a maximum (>200% of baseline) after 60 min. (Fig. 3D). In contrast, cPMN showed a different exocytosis behavior. CD66b surface expression was increased in both CCR5^−^ and CCR5^+^ cPMN only by ionomycin but not by fMLP. Moreover, the degree of ionomycin-induced CD66b upregulation was lower in cPMN compared to rPMN. Importantly, the overall increase of CD66b on CCR5^+^ cPMN was weaker compared to CCR5^−^ cPMN.

To examine phagocytosis, we incubated rPMN or cPMN with GFP-expressing *E. coli* for 15 minutes at 37°C, which led to cytochalasin D-sensitive phagocytosis (Supplementary Figure 2). Thereafter, cells were washed to remove free bacteria and analyzed using flow cytometry (Fig. 3E). 28±5% rPMN and 25±3% of CCR5^−^ cPMN, compared to 13±3% CCR5^+^ cPMN, phagocytosed at least one *E. coli* (Fig. 3F). Thus, cytokine-treatment *per se* did not interfere with exocytosis and phagocytosis of PMN, but a strong and significant reduction of these antibacterial properties could be observed for CCR5^+^ cPMN.

### CCR5^+^ cPMN are prone to spontaneous NET formation

Next, we analyzed the NETotic behavior of CCR5-defined populations of cPMN. CCR5^+^ cPMN were positively selected using magnetic beads, adhered on slides and NETosis was analyzed by confocal microscope as described (Fig. 4A) (*30, 31*). NET formation represents a multistep process including chromatin and nuclei reorganization, accompanied with a rounding and swelling of the nuclei eventually leading to chromatin expulsion (*30*). Thus, while the nuclei of mature PMN consist of 2-5 lobes, they de-lobe during NETing and appear with a spherical shape which may result in de-nucleated cytoplasts (*32, 33*) (Supplementary Figure 3A). In our experiments, CCR5^−^ cPMN showed mainly lobed nuclei (Fig. 4B). In contrast, CCR5^+^ cPMN showed a heterogeneous shape of their nuclei, including lobed nuclei (39% of the cells), round nuclei (35% of the cells), NET-shaped nuclei (8% of the cells) or denucleated cells (18% of the cells) which is related to NETosis (*32, 33*). Overview images confirmed ongoing NETosis of CCR5^+^ cPMN (Supplementary Figure 3B) and, thus, these data clearly indicate a pro-NETotic phenotype of CCR5^+^ cPMN.

**Figure 4.**
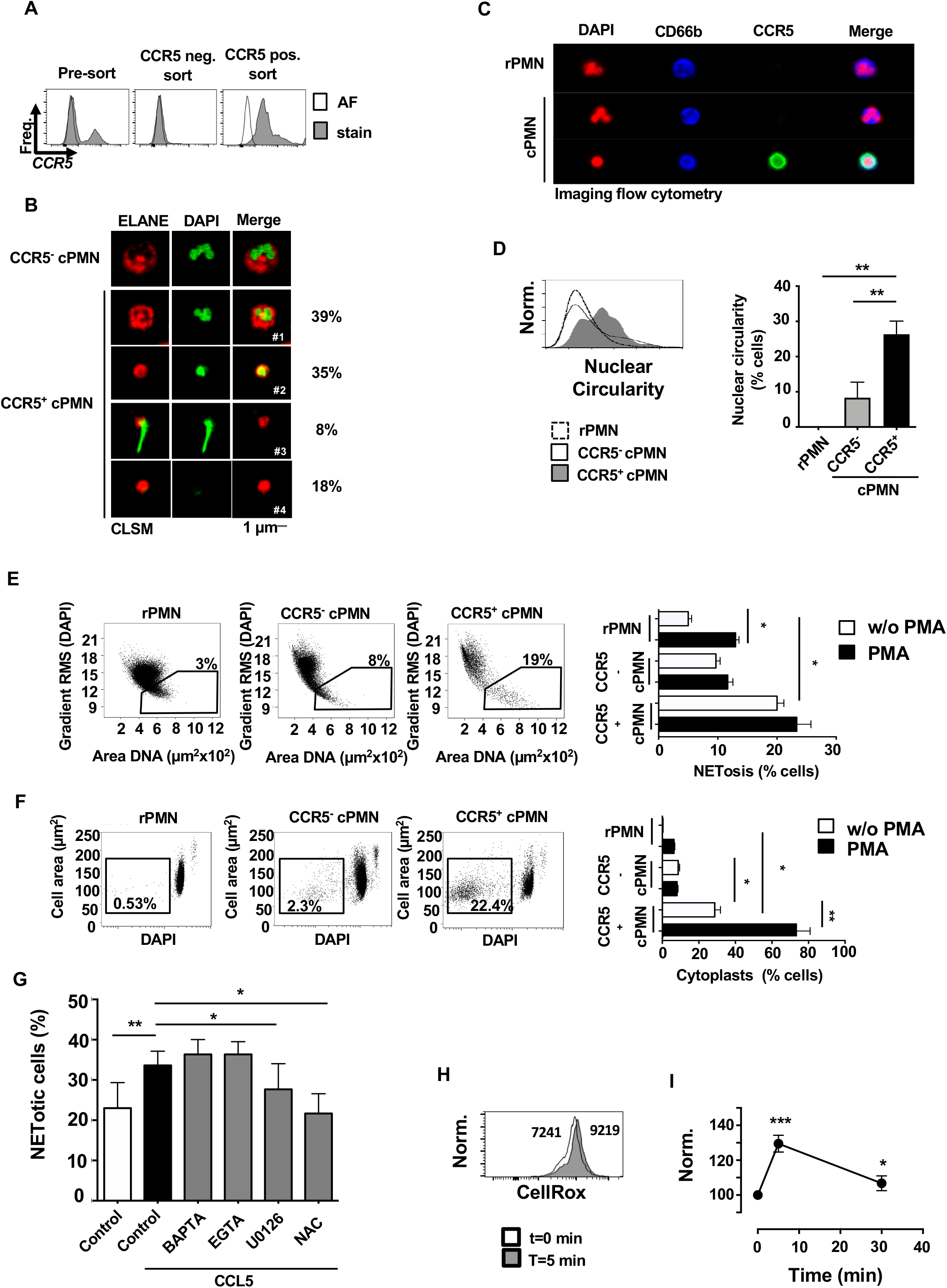
CCL5 induced ROS production accelerating NETosis of pro-NETotic CCR5^+^ cPMN. **A)** Flow cytometric analysis of cPMN before magnet-bead sorting, and the negatively sorted CCR5^−^ cPMN and positively sorted CCR5^+^ cPMN. The respective autofluorescence is shown as an open histogram. The figure is representative for 3 independent experiments. **B)** Microscopic confocal laser-scanned images of sorted CCR5^−^ cPMN (upper panel) and purified CCR5+ cPMN (lower panels). While CCR5^−^ cPMN mostly show a lobed nucleus, different nuclear morphologies can be seen in CCR5^+^ cPMN. The percentage of each morphology is shown on the right side and represents 3 independent experiments. **C)** Nuclear morphology was analyzed by imaging flow cytometry. PMN were fixed and stained with fluorescently labelled antibodies against CD66b (blue) and CCR5 (green) as well as with DAPI (red) to visualize nuclei. The images are representative for 6 independent experiments in which 25,000 cells were acquired. **D)** Nuclear circularity was quantified for rPMN (dotted line) and CCR5^−^ cPMN (solid line) and CCR5^+^ cPMN (grey histogram) by imaging flow cytometry. Data are representative for 6 independent experiments. A quantification is shown in the graph on the right (n=6; SEM, ANOVA, **p<0.01) **E)** To quantify NETosis, the chromatin-pattern was analyzed by imaging flow cytometry. A NETotic phenotype is present if the DNA-sharpness (y-axis) is small and at the same time the area on which the chromatin is distributed increases (x-axis). Shown are percent NETotic PMN in rPMN (left dot plot), CCR5^−^ cPMN (middle dot plot) and CCR5^+^ cPMN (right dot plot). The graph on the right show the mean percentages of NETotic PMN either unstimulated or after PMA-treated for 45 min (n=3; SEM; ANOVA, *p<0.05). **F)** Cytoplasts can be identified due to their depleted DNA content as measured by the intensity of DAPI-derived fluorescence of the cells. The sample dot plots and the quantification is analogous to the presentation in C (n=3; SEM; ANOVA, *p<0.05). **G)** cPMN incubated with CCL5 alone or together with the indicated reagents and incubated for 2h. The nuclear morphology was determined by imaging flow cytometry as described above (n=3; SEM; ANOVA, *p<0.05; **p<0.01). **H)** To determine the intracellular ROS level, cPMN were loaded with CellROX^®^ green and stimulated with CCL5 for 5 min. The flow cytometric histogram shows CellROX^®^ green in unstimulated cells (t=0 min, open histogram) and CCL5-treated cells (t=5 min, closed histogram). The numbers indicate the geo-mean values, obtained from the histograms in log-scale. The figure is representative for three independent experiments. **I)** The changes in ROS level after CCL5 treatment are shown as kinetics. Values are presented relative to unstimulated cells, which were set to 100% (n=3; SEM; ANOVA, *p<0.05; ***p<0.001)

To rule out an effect of positive selection on NETosis, we also measured NETosis by imaging flow cytometry without prior isolation. These analyses confirmed that rPMN and CCR5^−^ cPMN had mainly lobed nuclei, while lobed and round nuclei occurred in CCR5^+^ cPMN (Fig. 4C and D). In addition, imaging flow cytometry allows the calculation of the chromatin swelling (*30, 34*). This analysis revealed that only a minor proportion of rPMN and CCR5^−^ cPMN (3% and 8%, respectively) had swollen nuclei (Fig. 4E, left and middle dot plot), while a higher fraction of CCR5^+^ cPMN (19%) showed this nuclear phenotype indicating ongoing NETosis (Fig. 4E, right dot plot). For comparison, a known inducer of NETosis was applied, i.e. the phorbol ester PMA, which increased the level of NETosis-based nuclear morphology in rPMN significantly and of CCR5^−^ cPMN as well as CCR5^+^ cPMN by trend. To substantiate the NETosis dynamics in CCR5^+^ cPMN, an end-stage NETosis feature, i.e. the occurrence of denucleated cytoplasts, was evaluated (*32, 33*). A small fraction of denucleated cytoplasts in rPMN (0.53%) and CCR5^−^ cPMN (2.3%), but a high number of CCR5^+^ cPMN cytoplasts (22.4%) was observed (Fig. 4F). Importantly, PMA-stimulation significantly increased the occurrence of CCR5^+^ cPMN cytoplasts.

Because CCL5 induced polarization of CCR5^+^ cPMN but not migration after 3h (see above), it was reasonable to assume that CCR5-triggering activated signaling cascades that led to enhanced NETosis. To investigate this, cPMN were incubated with CCL5 for 3h and NETosis was investigated by evaluating nuclear morphology. Indeed, NETosis was again significantly increased by CCL5 in CCR5^+^ cPMN (Fig. 4G). This increase in NETosis was independently attenuated by the MEK inhibitor U0126 and the reducing agent/ROS antagonist N-acetyl-cysteine (NAC), but not by the calcium chelators BATA-AM or EGTA, suggesting that ROS and MEK-dependent increased ROS levels are likely to be responsible for CCL5-dependent increased NETosis (*35*). Indeed, CCL5 led to ROS production in CCR5^+^ cPMN (Fig. 4H), which was strongest after 5 min and deceased thereafter (Fig. 4I). Together, these results demonstrate that CCR5^+^ cPMN are prone to spontaneous NET formation and that CCR5 triggering amplifies NETosis.

### Increased expression and nuclear localization of ELANE in CCR5+ cPMN

We next investigated why CCR5^+^ cPMN *per se* exhibited a pro-NETotic phenotype. NETosis is a highly regulated process in which signaling cascades induce relocation of ELANE from azurophilic granules to the nuclei, where it participates in NETosis-related chromatin decondensation. Intracellular flow cytometry revealed a significant increase in ELANE expression in CCR5^+^ cPMN compared to both CCR5^−^ cPMN and rPMN (Fig. 5A and B). Imaging flow cytometry disclosed that ELANE was mainly localized in the cytoplasm in CCR5^−^ cPMN (Fig. 5C, upper panel). In contrast, ELANE displayed a nuclear localization in CCR5^+^ cPMN (Fig. 5C, lower panel, yellow color). A rescaled ELANE/DAPI Pearson correlation coefficient confirmed that ELANE was mainly in the cytoplasm of CCR5^−^ cPMN, and a nuclear translocation of ELANE was observed in 38.7±4.3% of CCR5^+^ cPMN (Fig. 5D and E). In conclusion, an increased expression and preferentially nuclear distribution of ELANE provide a mechanistic basis for the increased NETosis in CCR5^+^ cPMN.

**Figure 5.**
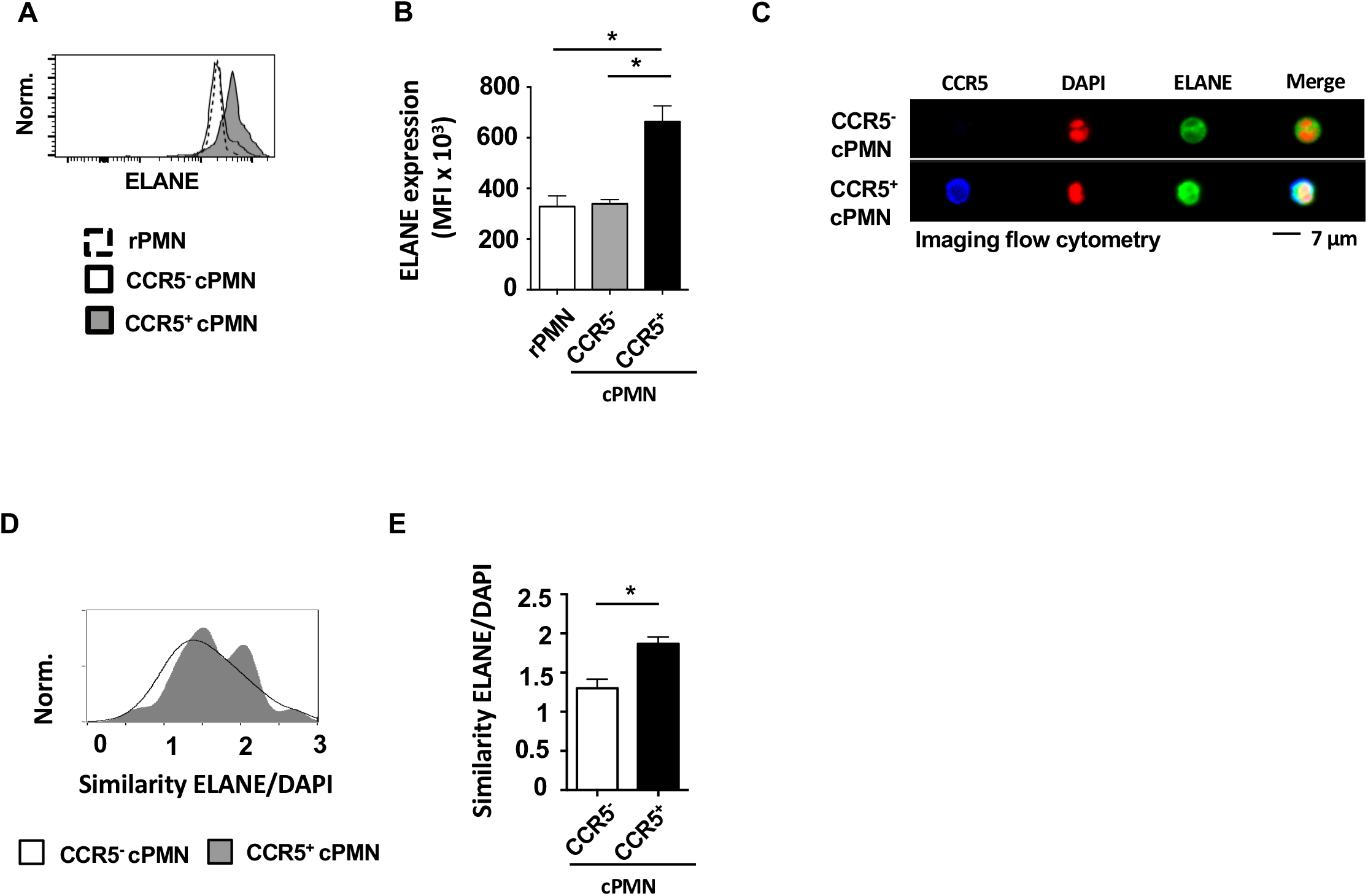
Increased ELANE expression in CCR5^+^ cPMN. **A-B)** The expression of ELANE was quantified by intracellular fluorescent immunolabelling. The grey histogram **(A)** shows the elevated ELANE level in CCR5^+^ PMN, while the ELANE expression of CCR5^−^ PMN (open histogram) and rPMN (dotted histogram) was on a comparable level. A statistical evaluation of three independent experiments is shown in **B** (n=3; SEM; ANOVA, *p<0.05). **C-E)** The subcellular localization of ELANE was analyzed using imaging flow cytometry. A co-localization of ELANE (green) and DAPI (red) appears in yellow in CCR5^+^ cPMN **(C)**. Nuclear localization of ELANE was quantified by calculation of the similarity score provided by imaging flow cytometry **(D)** and statistics were calculated for three independent experiments **(E)** (n=3; SEM; t-test, *p<0.05).

### Dichotomic function of soluble (sTNF) and membrane bound TNF (mTNF) on NETosis of CCR5^+^ cPMN

Since stimulation of PMN with GM-CSF plus IFNγ induced transcription of *TNF*, we speculated that IFNγ could induce TNF expression in cPMN which then could regulate ELANE expression and NETosis in CCR5^+^ cPMN in an para- or autocrine manner. By intracellular flow cytometry, TNF expression could already be detected at low levels in rPMN but increased in cPMN, especially in CCR5^+^ cPMN (Fig. 6A, left graph). The Tumor necrosis factor receptor 1 (TNFR1) was highly expressed on CCR5^−^ cPMN and dimly on CCR5^+^ cPMN or rPMN. Tumor necrosis factor receptor 2 (TNFR2) showed the opposite pattern, with high expression on CCR5^+^ but not CCR5^−^ cPMN (Fig. 6A, middle and right graph).

**Figure 6.**
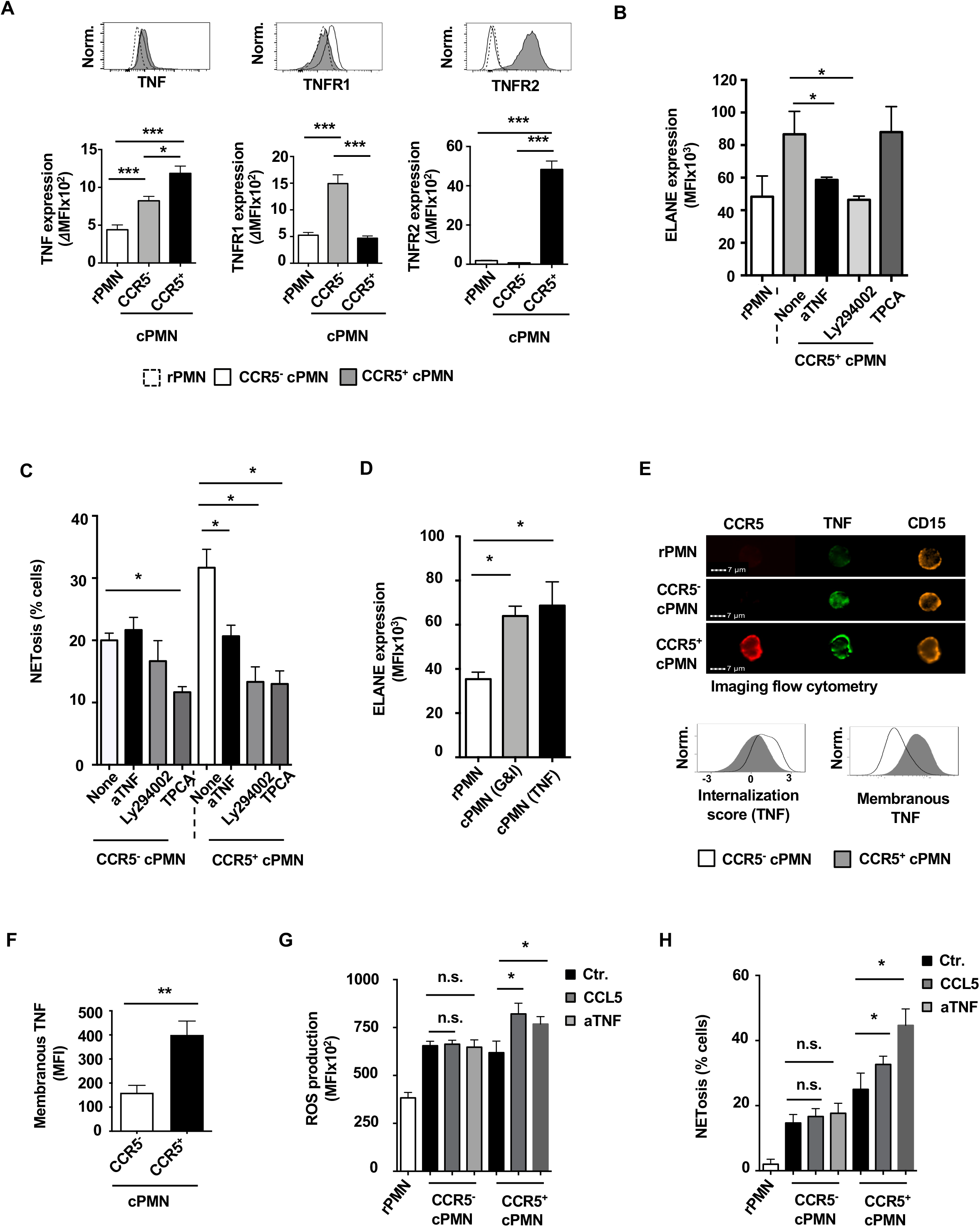
Membrane-bound TNF α mediates ELANE expression via TNFR2 and PI3K. **A)** Intracellular (TNFα) and surface (TNFR1, TNFR2) expression of TNFα and its receptors was analyzed on rPMN and cPMN by flow cytometry. The respective autofluorescence was subtracted from the respective MFI values to obtain the ΔMFI. The quantification shown below includes three independent experiments (n=3, SEM, ANOVA, *p<0.05; ***p<0.001). **B)** ELANE upregulation was measured in rPMN and CCR5^+^ cPMN. During the cytokine-dependent diversification process, cells were incubated with or without the PI3K inhibitor Ly294003 (Ly), an anti-TNF antibody (aTNF) or the NF-kB inhibitor TPCA (n=3; SEM; ANOVA, **p<0.01) **C)** PMN were incubated with either Ly294002, aTNF or TPCA during the diversification process. Subsequently, NETosis was detected by imaging flow cytometry (n=3, SEM, ANOVA, *p<0.05). **D)** PMN were incubated with GM-CSF plus IFNγ or TNF for two days and ELANE expression was determined by flow cytometry cytometry (n=4, SEM, ANOVA, *p<0.05). **E)** To investigate the localization of TNF in cPMN, cells were fixed, permeabilized, and treated with fluorescently labeled antibodies against TNF. Subsequently, the cells were analyzed by imaging flow cytometry. Typical images of rPMN, CCR5^−^ and CCR5^+^ cPMN are shown in the upper panel. In the lower part, the left histogram shows the internalization score of TNF and the right part shows the MFI of TNF on the membrane. Due to low expression, rPMN have not been evaluated here. The figure is representative of 3 experiments with different blood donors. **F)** Membrane-bound TNF was detected using fluorescently labelled antibodies that were applied to unpermeabilized cPMN (surface stain). The bar graph shows the statistical evaluation of three independent experiments (n=3, SEM, t-test, *p<0.05). **G)** After incubation with GM-CSF plus IFNγ for two days, the resulting cPMN were labeled with CellROX and then treated with CCL5 or antibodies directed against TNF (aTNF) for 5 minutes. ROS content was determined by flow cytometry (n=3, SEM, ANOVA, *p<0.05). **H)** After cytokine-induced diversification, cells were incubated with CCL5 or aTNFa for 2 hours. Subsequently, NETosis was measured by imaging flow cytometry (n=3, SEM, ANOVA, *p<0.05).

We next analyzed signaling pathways downstream of TNF. PI3Kinase inhibition but not IkB kinase β inhibition interfered with ELANE upregulation in CCR5^+^ cPMN (Fig. 6B). Notably, inhibition of PMN-secreted TNF by a blocking antibody also led to decreased ELANE expression in CCR5^+^ cPMN (Fig. 6B). Accordingly, application of anti-TNF or PI3Kinase inhibition during the diversification process reduced NETosis only in CCR5^+^ cPMN, but not in CCR5^−^ cPMN (Fig. 6C). In contrast, NF-kB inhibition reduced spontaneous NETosis in both CCR5^−^ cPMN and CCR5^+^ cPMN, suggesting that a process downstream of TNF is involved in ELANE expression. Indeed, stimulation of rPMN with recombinant human TNF (rhTNF) was sufficient to induce ELANE expression (Fig. 6D).

Since the two TNF receptors differ in their ability to recognize soluble versus membrane bound TNF, we examined the distribution of TNF in cPMN. Imaging flow cytometry revealed that the majority of cellular TNF was membrane-bound on CCR5^+^ cPMN, while it resided intracellularly in CCR5^−^ cPMN (Fig. 6E). mTNFα abundance was significantly higher on CCR5^+^ cPMN when compared to CCR5^−^ cPMN, suggesting that TNF-mediated outside-in signaling may be possible in CCR5^+^ cPMN (Fig. 6F).

To test this hypothesis, we applied anti-TNF antibodies to already diversified CCR5^+^ cPMN expressing mTNF and TNFR2. Flow cytometry revealed that cPMN contained higher ROS levels compared to rPMN. Anti-TNF treatment induced an additional increase in intracellular ROS, and a similar increase was found after addition of the CCR5 ligand CCL5 (Fig. 6G). Concordantly, NETosis was increased by anti-TNF treatment of cPMN (Fig. 6H). Thus, anti-TNF decreases ELANE expression and spontaneous NETosis when applied during the diversification of PMN but increases ROS-dependent NETosis when cPMN are already diversified (compare Model in Fig. 8) (*36*).

### CCR5^+^ PMN are abundant in the mucosa of patients with ulcerative colitis

PMN are associated with intestinal inflammation in inflammatory bowel disease, a disease group in which TNF blockade is often used successfully for treatment (*37, 38*). To determine whether increased CCR5 expression on PMN is detectable in IBD patients, sections of inflamed colonic tissue specimens of UC and CD patients and healthy controls (HC) were analyzed by immunohistochemistry. Azurocidin was used to visualize PMN (Fig. 7A, green). As expected, the frequency of PMN was higher in the lamina propria of CD and UC patients compared to controls. Notably, CCR5 colocalized with azurocidin and CCR5 expression on PMN could only be observed in the lamina propria of UC patients, but not in the colon of CD patients or healthy controls (Fig. 7A, red, quantification in Fig. 7B). In crypt abscesses, which are histologically defined by PMN exudates, the frequency of CCR5 expressing PMN was again significantly increased.

**Figure 7.**
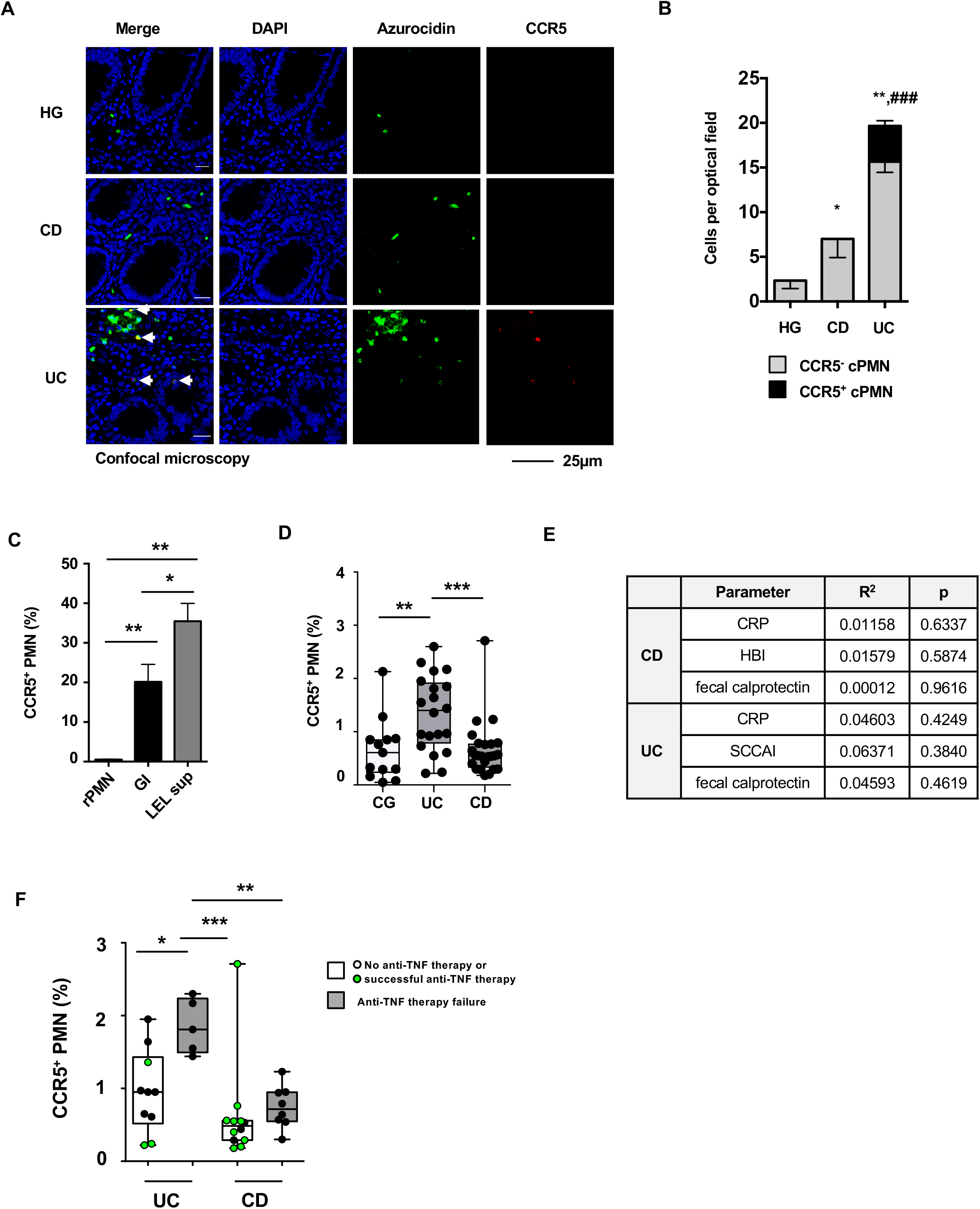
Frequency of CCR5^+^ cPMN ulcerative colitis patients. **A)** Tissue sections of mucosal specimens (CG, UC and CD) were subject to immunofluorescence staining of azurocidine (green) and CCR5 (red). Nuclei were counterstained with DAPI (blue). The arrows indicate CCR5^+^ cPMN in colonic mucosa of UC patients. Shown are the results from one representative experiment out of three independent experiments per condition. **B)** The bar graph shows the quantification of total PMN (grey) and CCR5^+^ cPMN (black) per optical field. In total 15 optical fields were quantified from three different patients (n=3; SEM; ANOVA, * p<0.05, ** p<0.01 of total PMN; ### p<0.01 of CCR5^+^ cPMN). **C)** Detection of the frequency of CCR5^+^ cPMN in rPMN or cells that were either stimulated with GM-CSF plus IFNγ (GI) or with LEL supernatants (n=3, SEM, ANOVA, *p<0.05; **p<0.01). **D)** The frequency of CCR5^+^ cPMN was quantified in the peripheral blood of a control group (CG) or UC and CD patients (SEM; Kruskal-Wallis, *p<0.01, ***p<0.001). **E)** A linear correlation of serum CRP level, the disease score (HBI: Harvey-Bradshaw Index or SCCAI: simple clinical colitis activity index) or fecal calprotectin level and the frequency of CCR5^+^ cPMN in the whole blood was calculated for the patients shown in E. **F)** The graph shows the percentage of CCR5^+^ cPMN in the whole blood of UC and CD patients that were naïve for anti-TNFα therapy (white column, black dots), were successfully treated with infliximab or adalimumab (white column, green dots) and patients in which the infliximab or adalimumab therapy failed (grey column, black dots).

We utilized the loss-of-epithelial-layer (LEL) system (*39*) to analyze whether the intestinal micro-millieu can induce diversification of human PMNs. In this explant model, the epithelial layer is detached from surgically obtained colon preparations by EDTA treatment, resulting in tissue inflammation and secretion of cytokines into the supernatant. We incubated human PMN from healthy donors with LEL supernatant and subsequently quantified the fraction of CCR5^+^ cPMN (Fig. 7C). These experiments demonstrated that LEL supernatant diversifies PMNs to CCR5^+^ cPMN, suggesting that CCR5^+^ cPMNs can develop locally in inflamed tissues.

We further examined whether CCR5 PMN could also be detected with surface staining of leukocytes in the peripheral blood of IBD patients. For this purpose, blood samples were collected from healthy controls and patients with UC and CD and flow cytometric analysis was performed without prior PMN isolation (Fig. 7D). Indeed, a significantly higher proportion of CCR5 cPMN was found in UC patients compared to HC and CD. The increased abundance of CCR5 PMN did not correlate with CRP level, disease score, or fecal calprotectin (Fig. 7E). However, we found a significant increase in the proportion of CCR5 cPMN in UC patients in whom anti-TNF therapy (infliximab or adalimumab) failed (Fig. 7F).

We conclude that soluble factors released from inflamed intestine in vitro induce a phenotype of CCR5 PMN which can also be found in inflamed mucosa in vivo and are detectable at increased frequency in the blood of patients who failed anti-TNF therapy.

## Discussion

PMN are not only bactericidal cells, but play a very important role in maintaining the inflammatory background of chronic inflammation. By producing extracellular traps and secreting of cytokines, PMN contribute to smoldering of inflammation and destruction of inflamed tissues. This function of PMN can be accompanied by PMN re-programming, i.e. a change in their phenotype and functionality. Here, we show that PMN can diversify into two distinct subgroups, i.e. CCR5^−^ (TNFR1^bright^CXCR4^bright^HLA-DR^dim^CD62L^dim^CD16^brigh^) and CCR5^+^ (TNFR2^bright^CXCR4^dim^HLA-DR^bright^CD62L^bright^CD16^dim^) cPMN. The latter subgroup showed decreased anti-bacterial armamentarium with the exception of NETosis, which was accelerated. CCR5^+^ cPMN were found in inflamed colon of UC patients. There, a NETosis-related release of ELANE might be an important trigger for the inflammation and tissue destruction in the gut of UC patients.

Expression of CCR5 on PMN is also evident in other contexts. A subgroup of PMN derived suppressor cells expressing CCR5 was described in mice (*40*). In this model, inhibiting CCR5 interfered with chemoattracting these cells into tumors and, thereby, interfered with their immuno-suppressive function. CCR5 expression on apoptotic PMN had anti-inflammatory functions by locally sequestering CCR5 ligands, a mechanism shown to take part in resolution of experimental peritonitis (*41*). This study supports our hypothesis that CCR5 transport to the cell surface represents a proximal priming event (Fig. 8A). Cytokine-mediated stimuli (e.g. GM-CSF), however, prevent apoptosis and prolong survival (*42*) as also evident by downregulation of *CASP8* gene in our study. Additional gene expression and TNF upregulation can be provoked by the IFNγ/ IFNR axis modulating PMN functionality. Thus, the sequence of events acting on PMN during the course of inflammation determine the fate of individual PMN.

**Figure 8.**
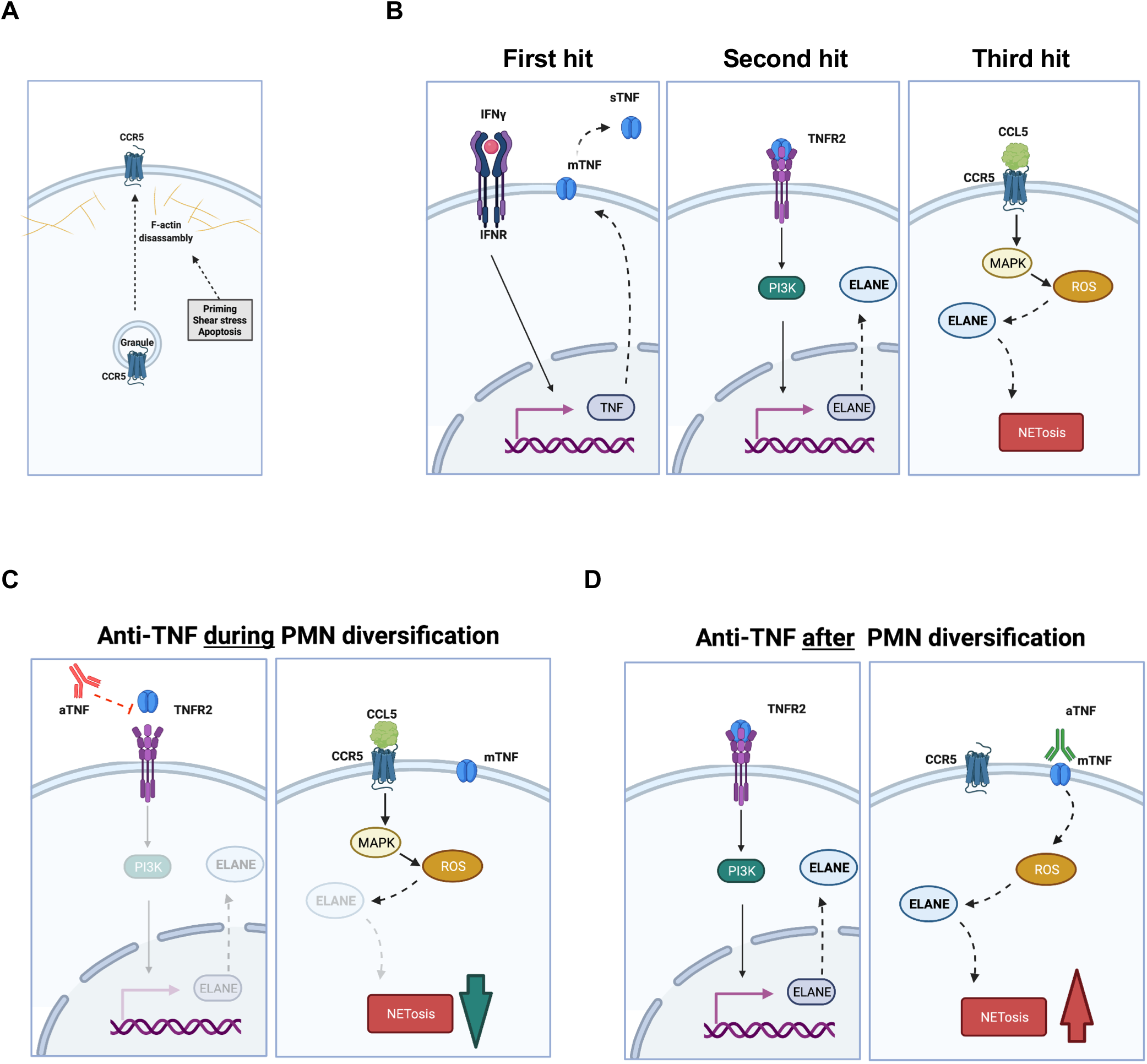
Three hit model of NETosis in inflammation. **A** Surface CCR5 expression is initiated on PMN by exocytosis after dissassambly of the F-actin barrie as part of a either priming, stress response or apoptosis. Without survival/ anti-apoptotic signals PMN undergo apoptosis at this stage. **B)** Enhanced CCR5 expression by *de novo* synthesis is induced and anti-apoptotic mechansism are initiated (not shown). In addition, a process comprising three hits drives PMN into NETosis. In the first hit, PMN are stimulated by IFNγ leading to an expression of TNF (left part). In the second hit, ELANE is overexpressed via a TNFR2 signaling pathway (middle part). TNF may have several sources. One is the CCR5^+^ cPMN by auto/paracrine stimulation via sTNF or mTNF. The third hit leads to activation of ELANE and induction of NETosis by ROS (right part). The latter can occur through stimulation of CCR5 with natural ligands. **C)** Blocking TNF antibodies can slow NETosis process by interfering with overexpression of ELANE downstrem of TNFR2. **D)** Antibodies to TNF can also induce an outside-in signal via mTNF. This produces ROS, which is an important mediator of NETosis. One function of ROS is to release ELANE from azurosomes, which accelerated NETosis.

Therefore, based on our data we propose the following model: PMN priming induces immediate transport of granule-stored CCR5 to the cell surface (Fig. 8A). A high deviation in the CCR5^+^ cPMN frequency occurred if PMN of the same donor were re-analyzed in otherwise independent experiments (data not shown) indicating that the observed diversification and development of CCR5^+^ cPMN is independent on epigenic mechanisms. Rather, this process is coupled to actin dynamics since perturbation of the actin cytoskeleton by CytoD increased CCR5 surface expression (*43–45*). At this stage, PMN are prone to apoptosis unless they receive survival signals. In the presence of cytokines, apoptosis is delayed, *de novo* synthesis reinforces CCR5 expression and a three hit sequence is initated in CCR5^+^ cPMN (Fig. 8B). In a first hit, IFNγ induces the expression of TNF. In the second hit, TNF binds to the TNFR2 leading to PI3K-dependent upregulation of ELANE expression and rendering (together with the disassembly of the actin cytoskeleton (*46*)) CCR5^+^ cPMN pro-NETotic. The third hit is mediated by increasing intracellular ROS levels. The amplification is due to ROS-dependent activation of ELANE (*47*) and is provoked by triggering of CCR5 dissociating CCR5 proximal signaling from CCR5-induced migration. This is in line with the reverting effect of ROS-scavenger NAC and with the upregulation of CYBB gene in cPMN. According to this three hit model, TNF plays a role for the development of CCR5^+^ cPMN as well as ELANE upregulation. It is known that the soluble moiety of TNF (sTNF) is released after proteolytic cleavage of the transmembrane isoform (mTNF) by TNF-converting enzyme (TACE; ADAM-17) (*48*). While TNFR1 binds soluble TNF, TNFR2-mediated signaling is also induced by sTNF and mTNF. Notably, soluble and membrane bound TNF can exert opposing biological effects and TNF directed antibodies can induce outside-in signaling or mTNF/TNFR2 crosslinking (*36, 49, 50*). The therapeutically used TNFR2-fusion protein (eternacept) does not bind to mTNF and can, thus, not induce outside-in-signaling. It was demonstrated that eternacept was not beneficial for treatment of IBD patients and patients with other autoimmune diseases and treated with eternacept had an increased risk of being diagnosed with IBD (*51, 52*). One possible explanation of this paradoxical effect is that eternacept shifts the balance of sTNF/TNFR1-toward mTNF/ TNFR2-signaling promoting the pro-NETotic state of CCR5^+^ cPMN. According to our model, anti-TNF antibodies can exert different biological effects on PMN, i.e. inhibiting ELANE expression and NETosis (Figure 8C) or induction of ROS and consequently accelerate NETosis (Figure 8D). Thus, besides the well described effects of anti-TNF therapy (*53–56*), its net result could be modulated by the PMN diversification state. This would explain why patients with higher CCR5^+^ cPMN frequency tend to be more prone to failure of anti-TNF therapy. Further prospective studies in colitis patients are needed to test this hypothesis.

Although there are a plethory of evidences showing that CCR5 is involved in chronic inflammatory diseases (reviewed in (*57*)) the current view on CCR5 in IBD is ambiguous. On the one hand, there is no gene polymorphism of CCR5 is associated with IBD (*58, 59*). On the other hand, the CCR5-ligands CCL3 (MIP-1α), CCL4 (MIP-1β), CCL5 (RANTES) and CCL8 (MCP-2) are expressed in the colon of IBD patients (*60*) and blockage of CCR5 ameliorated DSS/TNBS-induced colitis in mice models (*61*). In addition, dominant inactive CCL5 reduced tissue damage in experimental rat models (*62*). One reason for such ambiguity is that the individual frequency of CCR5^+^ cPMN in patients was not considered in previous studies and CCR5 detection in tissues has certain drawbacks: A plethora of spatial distributed CCR5 conformations and CCR5 oligomerization exsist (*63, 64*) and CCR5-specific antibodies display varying conformation-dependent affinities (*65*). This might also explain why PMN lacking CCR5 expression were observed in our gut tissue samples; antigen retrieval of FFPE-embedded tissue samples might lead to conformation-restricted epitope liberation. Nevertheless, infiltration of PMN into colonic mucosa and NETosis is considered a hallmark of UC (*66–70*), and PMN activity is substantially enhanced in the colon of active UC patients (*71*). Very little is known about the occurrence and function of PMN subgroups. The here described CCR5^+^ cPMN have pro-inflammatory properties. By producing TNF and an increased tendency to undergo NETosis, these cells likely contribute to disease deterioration rather than improvement. Since NET associated ELANE was shown to have higher proteolytic activity on the extracellular matrix compared to unbound ELANE (*72*), the NETosis-associated release of ELANE could be an important step for exacerbating colitis. In line with this assumption, fecal ELANE is a predictive clinical marker for activity of UC in patients (*73*).

PMN heterogeneity represents a still underestimated factor influencing the pathogenesis of inflammatory diseases. It can have different sources, and whether distinct subsets of PMN exist or if they represent different states along a continuous distribution is an area of active research (*12, 74, 75*). It was recently shown that PMN can be represented along a continuous, chronologically ordered spectrum, termed neutrotime, from which they can deviate to reach different polarization states as a function of tissue, stimulus and timepoint (*76*). Since inflamed PMN were mostly advanced along neutrotime, CCR5^+^ cPMN might develop from cells in late neutrotime. Deviation from neutrotime can occur at different stages, and the conditions in the immediate mirco-milleu as well as the sequence of stimuli determine PMN plasticity in inflammation. Thus, different flavours of CCR5^+^ cPMN might exsist and whether the here described CCR5^+^ cPMN and CCR5^−^ cPMN are temporally related activation states or divergent, independent subgroups originating from a common sequence and their relationship to other PMN described subgroups (*77*) remains to be determined.

Taken together, the here described biological properties of CCR5^+^ cPMN and their occurrence in UC-dependent cryptitis could represent another important piece of the puzzle for understanding UC pathogenesis and emphazises the need of standardized PMN phenotyping in inflammatory diseases.

## Material and Methods

### Antibodies and Reagents

The following primary antibodies were employed: CCR5-PE (Miltenyi, Bergisch Gladbach, Germany); CCR5 unlabeled; CCR1-APC-Fire 750, CXCR1-APC, CCR4-BV605, CD66b-PerCP-Cy5.5, CD15-AF594, CD62L-FITC, CD49d-APC, CD16-APC-Cy7, HLA-DR-AF700, TNFα-PE-Dazzle594, TNFR1-APC, TNFR2-PE-Cy7 (Biolegend, San Diego, USA); CXCR3-APC, CXCR3-PE-Cy5, CXCR4-PE-Cy5, CCR3-PE CCR3-PE-CF594, CD15-BV605, CD49d-PE-Cy5 (BD Bioscience, Heidelberg, Germany); ELANE (MerckMillipore, Darmstadt, Germany); CXCR4-FITC (R&D Systems, Minneapolis, USA); TNFα (purified), CCR5 (purified) (R&D Systems, Minneapolis, USA); azurocidin (Abcam, Cambridge, UK); Anti-mouse-Cy5 (Dianova, Hamburg, Germany). DAPI, Phalloidin, ionomycin, paraformaldehyde, PMA and CytoD were from MerckMillipore (Darmstadt, Germany), TPCA from Abcam (Cambridge, UK), GM-CSF and CCL5 from R&D Systems (Minneapolis, USA), IFNγ from Biolegend (San Diego, USA).

### Isolation of human peripheral blood neutrophils (PMN)

PMN were purified from 30-90ml peripheral blood of healthy donors. The isolation of PMN was performed through density centrifugation. Briefly, for each preparation 30ml of whole blood was added to 20ml of PolymorphPrep™ and centrifuged at 535g for 35 min. The PMN layer was collected, washed with 0.45% sodium chloride solution in a 1:1 ratio and centrifuged at 443g for 10 min. To remove erythrocyte contamination, the pellet was mixed with 20ml of 0.2% sodium chloride solution. The resulting hemolysis then was stopped after 10 seconds through adding 20ml of 1.6% sodium chloride solution. After washing twice, the pellet was resuspended in RPMI + 10% FCS. For mRNA analysis, a second bead-based purification step was included according to the manufacturer’s instructions (EasySep^TM^ Human Isolation Kit from STEMCell Technologies, Cologne, Germany). While density centrifugation led to 90% pure PMN, the two-step isolation protocol allowed to reach 99.7% PMN purity. After purification, cells were seeded with a mixture of GM-CSF (100 U/ml) plus IFNγ (10 ng/ml) in 6-well-plates (5 ml per well) and incubated at 37°C and 5% CO_2_ for 48 hours.

For bead-based isolation of human CCR5^+^ and CCR5^−^ cPMN were sorted using anti-APC MojoSort™ nanobeads (Biolegend, San Diego, USA) according to the manufacturer’s recommendations. Briefly, PMN were stimulated with GM-CSF (100 U/ml) plus IFNγ (10 ng/ml) for one day. 2×10^6^ cells were collected and labeled with APC-labelled CCR5 antibodies (Miltenyi, Bergisch Gladbach, Germany) in RPMI supplemented with 10% FSC, 2 mM EDTA and 10 mM HEPES for 20 min. After washing 12.5μl anti-APC MojoSort™ nanobeads and 1 ml RPMI supplemented with 10% FSC, 2 mM EDTA and 10 mM HEPES were added. After 20 min. incubation time, cells were isolated using a magnet.

### Flow cytometry

PMN were fixed with 1.5% PFA for 10 min followed by one washing step with FACSwash buffer (PBS, 10g/l BSA, 0.1% NaN_3_ – for intracellular staining 0.1% saponin was added). All washing steps were completed by centrifugation at 300 g for 6 min. The PMN pellet was then incubated light-protected with 50 μl of FACSwash-antibody-mix for 20 minutes. Thereafter, excess antibodies were removed by washing with FACSwash buffer. For whole blood staining, 200 μl of heparinized whole blood were incubated with 50 μl antibody-mix for 30 min protected from light at 4°C. Red blood cells were removed by adding 2 ml of BD FACS Lysing Solution (BD Bioscience, Heidelberg, Germany) for 10 min at room temperature. The cells were then washed twice with FACSwash buffer (PBS, 10g/l BSA, 0.1% NaN_3_, 0.1% saponin). Cells were measured using an LSRII flow cytometer (BD Bioscience, Heidelberg, Germany) and analyzed using FACSDiva™ software version 8 and FlowJo™ version 10.6.0 (BD Bioscience, Heidelberg, Germany). To identify PMN, cells were gated according to the FSC/SSC profile combined with CD66b expression.

### Imaging flow cytometry

For imaging flow cytometry, samples were prepared as described for conventional flow cytometry (see above), and cells were finally resuspended in 50μl PBS/ 5%BSA. Thereafter, 25,000 cells were acquired using an ImageStream IsX MkII and Inspire software (Amnis, Seattle, USA). Data analysis was performed with IDEAS 6.2.188.0 (Amnis, Seattle, USA) (*78, 79*). Briefly, cells were gated according to the cell size (area brightfield) and side scatter signal as well as their CCR5-expression. Cell polarization was quantified according to the aspect ratio (AR) of the phalloidin signal. Polarized cells were considered as such, if the AR was below 0.85. For assessing the nuclear morphology on the single cell level, an DAPI-dependent threshold mask was applied (M07, Ch07, 40; (for details about masking see (*78, 80, 81*))), and the distances of the mask center to the boundaries were measured (circularity feature). Higher circularity values indicated rounder shapes of the nuclei, whereas low values indicated lobular nuclei. During NETosis the nuclear membrane dissolves and the chromatin begins to swell. Microscopically, chromatin swelling can be determined by a decrease in the contrast of nuclear images combined with an increase of the area covered by the chromatin (*34*). To quantify these morphological characteristics of the chromatin, we calculated the root mean square for image sharpness, i.e. the “Gradient RMS” feature, which correlates with the contrast of the nuclear images and was calculated by imaging flow cytometry and IDEAS software. In addition, the area of the nuclei in metric numbers (μm^2^) was calculated.

### Phagocytosis assay

Ampicillin-resistant, GFP-expressing *Escherichia coli* were grown overnight in lysogeny broth (LB medium) at 37°C with continuous rotation at 230 rpm. After 12 hours, the number of bacteria was assessed using turbidity measurement and adjusted to 10^7^ bacteria/ml in RPMI + 10% FCS. PMN were prepared in aliquots of 1 ml (RPMI + 10% FCS) containing 1×10^6^ cells. To each PMN sample, 50 μl *E. coli* were added to achieve a PMN-bacteria-ratio of 1:10 or 1:12, respectively. The samples were incubated at 37°C for 20 min and washed once with RPMI + 10% FCS and once with PBS. Cells were fixed and stained with fluorochrome labelled antibodies as indicated and measured immediately with conventional or imaging flow cytometry.

### Loss-of-epithelial layer model

Mucosal specimens were washed extensively in RPMI 1640 (Life Technologies, Paisley, UK) containing antibiotics prior to removing the mucus layer by dithiothreitol (DTT) treatment (1 mM, 10 min; Sigma, St. Louis, MO, Germany). Subsequently, punches of defined surface area were prepared and denuded of epithelial cells by exposure to 0.7 mM EDTA (Sigma) in HBSS (without Ca^2+^ and Mg^2+^; Life Technologies) at 37°C in a shaking water bath for 30 min (50 ml EDTA/HBSS per punch) (*82*). This incubation was repeated three times with two washing steps (10 min in HBSS/antibiotics) after each incubation period. Subsequently, mucosal specimens were cultured in a defined volume of medium (*39, 82*) for 12h. After 12h of organ culture, the supernatant was harvested, centrifuged and used for incubation of PMN.

### Immunohistochemical staining

Biopsies of colonic mucosa were taken from patients diagnosed with active state of inflammatory bowel disease (IBD), i.e. ulcerative colitis (UC) or Crohn`s disease (CD). Immunofluorescence staining was performed using 2 μm sections of formalin-fixed, paraffin embedded (FFPE) human colon tissue previously fixed in 4% PBS-buffered methanol-free formaldehyde (Brenzinger, Walldorf, Germany) for 2h. After deparaffinization, rehydration and heat-mediated antigen retrieval blocking was performed by incubation with an antibody diluent. Immunofluorescence staining was performed using OPAL™ technology (PerkinElmer, Waltham, MA, USA). Procedures were performed according to the manufacturer’s instructions.

### Structured-illumination microscopy

Cells were re-suspended in RPMI 1640 without FCS at a density of 1×10^6^ cells/ml. These cells were seeded on poly-D-lysine (final conc.: 0.01mg/ml, Merck, Darmstadt, Germany) coated coverslips (Ø10mm, 1.5H high-precision, Marienfeld, Laud Königshofen, Germany) and incubated for 15min at 37°C. Cells were fixed using 1.5% PFA and stained as indicated in the figures. Coverslips were finally washed three times with PBS containing 1% BSA, 0.1% saponin and 0.07% NaN_3_ and finally with deionized water. Coverslips were then mounted on a microscopy slide (ProLong Gold, Thermo Fisher Scientific). 3D-SIM images were taken at latest two days after sample preparation with a Nikon N-SIM microscope (Nikon, Tokio, Japan) using a 100x objective (SR Apo TIRF100x NA 1.49). The reconstruction parameters were set as following: Illumination Modulation Contrast: 0.08; High resolution noise suppression: 0.8; Out of focus blur suppression: 0.8. If Illumination Modulation Contrast was set to auto-detection, the reconstruction quality was calculated between 7-8 (scale 1-10). Image analysis and lookup table adjustments were performed using NIS-Elements V 4.30 (Nikon).

### Confocal laserscan microscopy (NETosis detection)

Immunofluorescence staining were essentially done as described (*31*). Briefly, PMN were plated on slides for 15 minutes. All following steps were performed at room temperature. Cells were washed with PBS and fixed in 1.5% paraformaldehyde in PBS overnight. Cells were permeabilized with 0.3% Triton X-100 and stained with DAPI. 3D-SIM images were taken within two days after sample preparation with a Nikon A1 microscope using a 20x objective.

### Expression analysis

Total RNA was isolated from rPMN and cPMN of three healthy blood donors. PMN were isolated by gradient centrifugation followed by magnetic bead-based separation. Total RNA was then purified from 10,000 cells using MagNA Pure (Promega, Mannheim, Germany) in the inhouse molecular immunodiagnostic facility according to the manufacturer’s instructions. All RNA samples were quantified by using Qubit RNA assay kit (Thermo Fisher Scientific, Waltham, Massachusetts), and RNA integrity was assessed using Agilent 2100 Bioanalyzer system. Gene expression was analyzed in collaboration with the NanoString Core facility of Heidelberg. Briefly, 25 μg total RNA (5 μL/sample) was mixed with nCounter® reporter CodeSet (3 μL) and nCounter® capture ProbeSet (2 μL) with hybridization buffer (5 μL) for an overnight hybridization reaction at 65 °C. The reaction was cooled down to 4 °C, the samples were purified and immobilised on a cartridge, and data was assessed on the nCounter SPRINT Profiler. The exported data was analysed using Nanostring nSolver 4.0. For downstream analysis, only genes with an expression value of ≥ 100 CodeSet counts in all three replicates of either rPMN or cPMN or both were considered. Differential expression analysis was performed using limma with P values adjusted for multiple comparisons according to Benjamini and Hochberg, corresponding to a false discovery rate (*83*). Genes with an expression of below this threshold were not further analysed (*84*). Differentially expressed gene sets were mapped onto gene interaction networks from STRING v.11 (*85*). STRING integrates protein-protein interactions from literature curation, computationally predicted interactions, and interactions transferred from model organisms based on orthology.

### Ethics statement

All human studies were approved by the ethics committee of the University of Heidelberg (ethical votes S-024/2003, S-285/2015, S-390/2015 and S-119/2017) and performed in accordance with the principles laid down in the Declaration of Helsinki. Written informed consent was obtained from the patients.

### Statistics

Statistical analysis was performed using GraphPad Prism V6 (La Jolla, CA, USA). For experimental data, means and standard errors of the means (SEM) are shown. Two groups are compared by two-sided t-tests and multiple comparisons by ANOVA and Fisher’s LSD test. Evaluation of PMN from the peripheral blood of IBD patients is depicted as box plot and analyzed by Kruskal-Wallis test.

## Acknowledgements

The authors thank Dr. Felix Lasitschka, Antje Heidtmann und Sabine Wendtrup for their help in tissue preparation and staining, Prof. Dr. Maria Hänsch and PD Dr. Luise Erpenbeck for critical reading of the manuscript draft, Ralph Röth and Heike Kuzan from the nCounter Core Facility Heidelberg for performing nCounter analyses and related services. There are no conflicts of interest.

