## Supplement Data for "Inflammation induces pro-NETotic neutrophils via TNFR2 signaling"

### Supplementary Data

Friederike Neuenfeldt[1,2*], Jan Christoph Schumacher[1,2*], Ricardo Grieshaber-Bouyer[3], Jüri Habicht[1,2], Jutta Schröder-Braunstein[2], Annika Gauss[4], Beate Niesler[5,6], Niko Heineken[1,2], Alexander Dalpke[7], Matthias M. Gaida[8], Thomas Giese[2], Stefan Meuer[2], Yvonne Samstag[1,2], Guido Wabnitz[1,2]

[1] Section Molecular Immunology, Heidelberg University, Im Neuenheimer Feld 305, D-69120 Heidelberg, Germany
[2] Institute of Immunology, Heidelberg University, Im Neuenheimer Feld 305, D-69120 Heidelberg, Germany
[3] Department of Medicine V, Hematology, Oncology and Rheumatology, Heidelberg University Hospital, Heidelberg, Germany
[4] Department of Gastroenterology and Hepatology, University Hospital Heidelberg, Im Neuenheimer Feld 410, D-69120 Heidelberg, Germany.
[5] Department of Human Molecular Genetics, Heidelberg University, Im Neuenheimer Feld 366, D-69120 Heidelberg, Germany.
[6] nCounter Core Facility, Department of Human Molecular Genetics, Heidelberg University, Im Neuenheimer Feld 366, D-69120 Heidelberg, Germany.
[7] Institute of Medical Microbiology and Hygiene, Technische Universität Dresden, Dresden, Germany.
[8] Institute of Pathology, University Medical Center Mainz, JGU-Mainz, 55131 Mainz, Germany.

[*] FN and JCS contributed equally to this manuscript.

Keywords: Neutrophils, Neutrophil diversification, Neutrophil subgroups, Ulcerative colitis, NETosis, TNF

**Supplementary Figure 1**

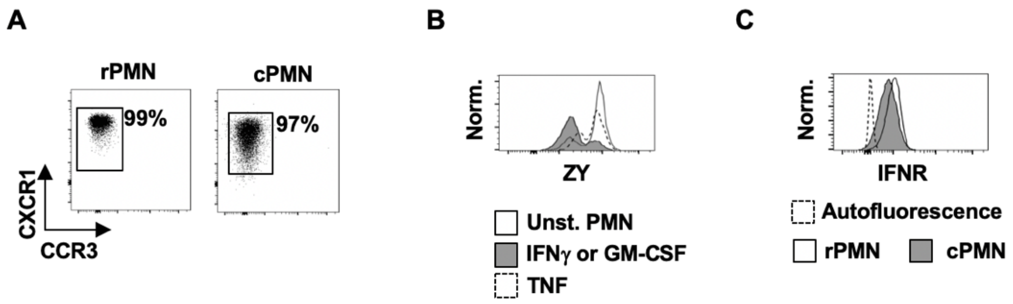


**Purity and IFNR expression of isolated human PMN**

**A)** PMN (rPMN and cPMN) were stained with fluorochrome labelled antibodies against CXCR1 and CCR3. The latter is expressed on eosinophils. Purification of human PMN resulted in one homogenous CXCR1$^{high}$CCR3$^{-}$ rPMN population. Stimulation of these cells with GM-CSF and IFNγ for two days (cPMN) did not induce CCR3 expression. The dot plots are representative for three independent experiments.

**B)** Primary human PMN were incubated in the presence of GM-CSF and IFNγ or TNF or left unstimulated for two days. Thereafter, cell viability was assessed using Zombie yellow (ZY).

**C)** The IFN receptor (IFNR) was expressed on all human rPMN and cPMN. The expression of IFNR was assessed by flow cytometry. The histogram is representative for three independent experiments with three different blood donors.

**Supplementary Figure 2**

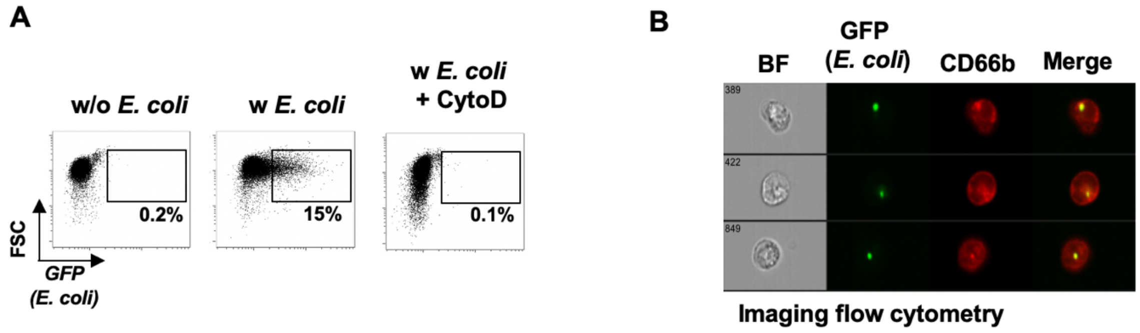


**Cytochalasin D inhibits phagocytosis of PMN**

**A)** Phagocytosis of *E. coli* by PMN was analyzed by imaging flow cytometry. PMN were co-cultured with (or without) GFP-expressing *E. coli* for 10 min. as indicated. As control, PMN were pre-incubated with CytoD (5µg/ ml) before *E. coli* were added. After PFA-fixation, the cells were analyzed by imaging flow cytometry and percentage of PMN that phagocytosed at least one *E. coli* was assessed.

**B)** Sample images of PMN that have phagocytosed *E. coli*. Data are representative for three independent experiments.

#### Supplementary Figure 3

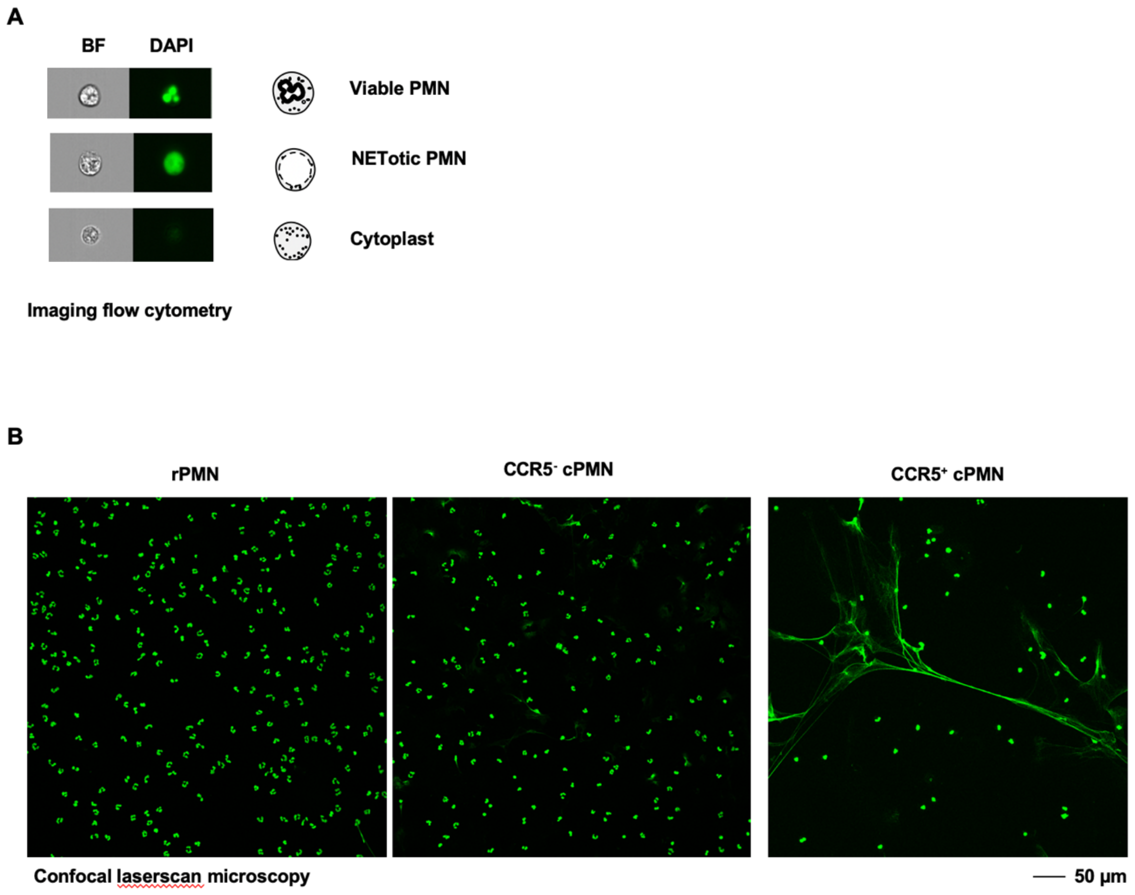


**A)** Imaging flow cytometry images of NETosis: Shown are representative brightfield (BF) and nuclear images (DAPI) as well as cartoons of a viable PMN (upper part), a NETotic PMN with decondensed DNA (middle part) and a denucleated cytoplast (lower part).

**B)** Confocal laserscan images of rPMN and purified CCR5+ and CCR5- cPMN. The figure shows DAPI stained cells that were acquired with an 20x objective (NA 0.8).
